# Microbial cross-feeding interactions reshape evolutionary trajectories of consumers by preserving motility

**DOI:** 10.64898/2026.05.25.727712

**Authors:** Thibault Rosazza, Zahraa Al-Tameemi, Alejandra Rodríguez-Verdugo

## Abstract

Cross-feeding interactions, in which a producer species release by-products that serve as resources for a consumer species, play an important role in shaping microbial community diversity. Producers create opportunities for consumers by supplying high-energy resources that are often scarce in the environment. However, they also exert strong top-down effects by releasing metabolites in pulses and generating spatial gradients of resource availability. How these spatiotemporal constraints shape consumer evolution remains poorly understood. To address this question, we used a two-species cross-feeding system in which *Acinetobacter johnsonii* excretes benzoate (a by-product of benzyl alcohol oxidation) into the external environment where it is consumed by *Pseudomonas putida*. To assess how the origin of benzoate (externally supplied or produced by cross-feeding) shapes consumer evolution, we evolved *P. putida* for 200 generations in monoculture or in co-culture with *A. johnsonii*. Populations evolved in monoculture exhibited improved growth relative to the ancestor, whereas populations evolved under cross-feeding showed little to no growth improvement.

Whole-genome sequencing revealed pervasive loss-of-function mutations in flagellar genes among populations evolved in monoculture, but not under cross-feeding conditions. High-throughput imaging assays showed that populations evolved under cross-feeding not only maintained but also enhanced functional motility. Competition experiments with single mutants revealed context-dependent fitness effects: loss-of-function mutations were highly beneficial when benzoate was externally supplied but deleterious when benzoate was supplied by *A. johnsonii*, highlighting the importance of motility in cross-feeding interactions. Together, our results show that resource origin fundamentally reshapes selective pressures and alters evolutionary outcomes in microbial communities.

**Significance Statement:** Cross-feeding interactions are pervasive in microbial communities, often arising from metabolite leakage into the environment. Our study focused on understanding how cross-feeding interactions shape the evolution of populations that cross-feed on these resources. Using a highly trackable cross-feeding system and linking genotypic to phenotypic changes, we showed that cross-feeding interactions reshape the evolutionary trajectories of consumer species. Notably, selection acts on flagellar genes, but their effects on function depend on the origin of resources. When resources are externally supplied, selection favors loss of flagellar motility, whereas when resources are generated through cross-feeding, selection not only maintains functional motility but also enhance it. These findings highlight that navigating the environment is essential for exploiting high-energy resources generated through cross-feeding.

## Introduction

Metabolic interactions are prevalent in microbial systems and underlie the functioning and stability of microbial communities (1, 2). Metabolic exchanges arise from the intrinsic propensity of microbial cells to leak metabolites due to overflow metabolism (3). Once in the external environment, these metabolites become resources for other species, i.e., cross-feeding interactions (4). Cross-feeding interactions are often unidirectional, in which one species (the producer) releases metabolites used by another (the consumer) without reciprocation. In unidirectional cross-feeding, the producer is minimally affected by the interaction, whereas the consumer gains a substantial advantage by accessing untapped and often high-energy resources. However, this opportunity to exploit high-energy resources comes with spatiotemporal constraints. First, resources produced by cross-feeding impose temporal constraints because they are often released in pulses rather than continuously. These pulses arise because intracellular metabolite concentrations must exceed a threshold to passively leak into the external environment (3). At the community level, the producer must reach a critical population density to exert a significant effect on other species (5). Second, resources generated through cross-feeding impose spatial constraints. Often, producers generate local metabolite gradients, especially in spatially structured environments, with concentrations highest near producer cells (6). Therefore, biogenic resources, defined here as resources generated by cross-feeding species, are highly dynamic and spatially structured, especially compared to homogenously distributed resources that are readily available in the environment, here called non-biogenic resources. Because biogenic and non-biogenic resources differ in their spatiotemporal properties, they are expected to impose distinct selective pressures on consumers. However, it remains unclear how consumer adaptation to biogenic resources differs from adaptation to non-biogenic resources.

To address this question, we leveraged a well-established synthetic microbial community composed of two bacterial species engaged in unidirectional cross-feeding (7–11). The producer, *Acinetobacter johnsonii,* degrades benzyl alcohol into benzoate, an intermediate product of benzyl alcohol oxidation. Benzoate accumulates intracellularly before leaking in small quantities into the extracellular environment, where it can subsequently be reutilized and fully degraded (7, 9). While *Pseudomonas putida* is unable to use benzyl alcohol, it cross-feeds on the extracellular benzoate secreted in pulses by *A. johnsonii*. This synthetic community provides a powerful model system for studying cross-feeding, as both species are genetically tractable, and their interactions can be manipulated by controlling the chemical composition of the culture medium.

Using this system, we conducted a fully factorial evolution experiment that manipulated both resource origin (biogenic versus non-biogenic) and resource concentration. This design enabled precise control over the extent to which *P. putida* depended on *A. johnsonii* for essential resources (Fig. 1A). Under obligate cross-feeding, *P. putida* relied entirely on biogenic benzoate, whereas under facultative cross-feeding, it had access to both biogenic and non-biogenic benzoate. In the absence of *A. johnsonii* (monocultures), *P. putida* used non-biogenic resources externally supplied.

**Figure 1.**
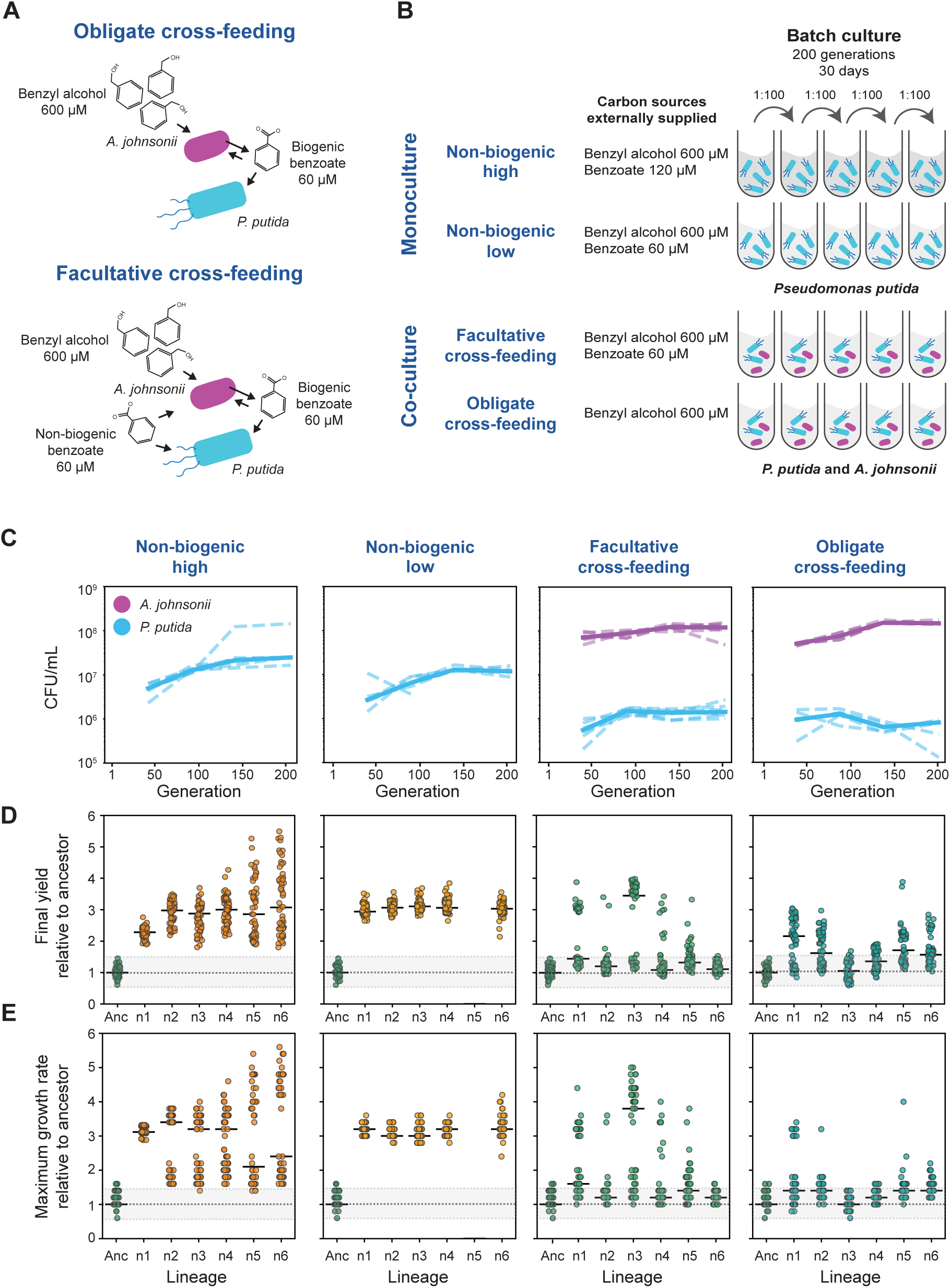
Unidirectional cross-feeding constrains improvement of growth-associated traits in *P. putida.* (*A*) Schematic representation of the obligate and facultative commensal cross-feeding interaction between *A. johnsonii* and *P. putida*. When benzyl alcohol is supplied as the sole carbon source, *P. putida* relies on the biogenic carbon released by *A. johnsonii* (obligate cross-feeding). Note that benzoate released by *A. johnsonii* can also be a carbon source for this species. When both benzyl alcohol and benzoate are supplied as sole carbon sources, *P. putida* can either use the non-biogenic carbon source directly or benefit from the biogenic carbon source released by *A. johnsonii* (facultative cross-feeding). (*B*) Schematic representation of the evolution experiment performed in this study. (*C*) Line plot representing the CFU/mL for *P. putida* (cyan) and *A. johnsonii* (magenta) of the four different evolved conditions at generations 50, 100, 150 and 200. The dark solid lines represent the median of 6 evolved lineages; dashed lines represent individual evolved lineages. (*D* and *E*) Scatterplot showing the yield (panel *D*) and maximum growth rate (panel *E*) relative to the ancestral population for 60 colonies from evolved *P. putida* populations across all replicates in each condition at generation 200. The gray area represents the zone of non-significant differences, and the black lines represent the median values for the 60 colonies tested.

We found that the consumer (*P. putida*) followed two evolutionary trajectories depending on the resource origin (i.e., biogenic vs non-biogenic). When supplied with low concentrations of non-biogenic resources, consumer populations adapted to nutrient starvation by enhancing growth-associated traits and, accumulating loss-of-function mutations in flagellar genes. In contrast, when equivalent concentrations of resources were produced by cross-feeding (i.e., biogenic resources), consumer populations accumulated a different set of mutations than in monoculture and not only retained but also enhanced motility. We further showed that the fitness effects of the flagellar mutations were context-dependent, suggesting that antagonistic pleiotropy underlies the observed differences in adaptive trajectories. Together, these findings demonstrate that the resource origin fundamentally reshapes selective pressures, with important consequences for the evolution of foraging traits such as motility.

## Results

### Cross-feeding interactions constrain evolution toward improved growth

To investigate how cross-feeding interactions influence the evolution of *P. putida*, we conducted a fully factorial evolution experiment (Fig. 1B). Under the obligate cross-feeding condition, benzyl alcohol was supplied as the sole carbon source, such that *P. putida* relied exclusively on benzoate produced by *A. johnsonii*. Under the facultative cross-feeding condition, both benzyl alcohol and benzoate were supplied, allowing *P. putida* to use biogenic and/or non-biogenic benzoate. In monoculture conditions, *P. putida* relied on externally supplied non-biogenic benzoate. Two benzoate concentrations (’low’ and ’high’) were used to match concentrations inferred for obligate and facultative cross-feeding, respectively, based on high-performance liquid chromatography measurements (9). We experimentally evolved six replicate populations under each condition for approximately 200 generations (30 days, Fig. 1B) and observed that all lineages persisted during the evolution experiment except for one replicate (n5) in the monoculture condition with low benzoate, which went extinct after ∼92 generations and was removed from further analysis (Fig. 1C).

To evaluate how cross-feeding interactions shape the phenotypic evolution of *P. putida*, we characterized the growth of the evolved populations and compared it with that of the ancestral population. We sampled 60 random clones from the ancestral and evolved populations (at generation 200) and measured growth curves in minimal media supplemented with benzoate as the sole carbon source. From these growth curves, we quantified four growth traits: final yield, maximum growth rate, half-saturation constant, and the maximum uptake rate.

In monoculture, *P. putida* populations evolved a higher final yield and maximum growth rate relative to the ancestor across all lineages (Fig. 1D-E and Fig. S1A-B), indicating that when evolving on a non-biogenic carbon source, *P. putida* populations adapted to the media conditions by enhancing growth traits. In contrast, populations evolving in co-culture exhibited fewer clones with enhanced growth. Most of the clones (ranging from 55% to 100% across all lineages) exhibited yields and growth rates similar to those observed in the ancestral population (Fig. 1D-E and Fig. S1A-B).

Furthermore, clones with improved growth from the facultative cross-feeding condition exhibited a higher yield compared to the obligate cross-feeding condition (average yield: 0.025 and 0.018 OD per mM for facultative and obligate cross-feeding, respectively; ANOVA, *p* = 1^e-7^). These results suggest that *P. putida*’s growth improvement is limited when evolving on a biogenic carbon source, and that the more *P. putida* is dependent on *A. johnsonii* for benzoate, the more its growth is constrained.

### Loss-of-function mutations in the flagellar operon occur less frequently under cross-feeding conditions

Next, we sought to link these previously quantified phenotypic observations in evolved *P. putida* populations with the acquisition of *de novo* mutations. To this end, we performed whole-population sequencing of evolved *P. putida* populations at 50, 100, 150, and 200 generations and quantified the frequency of *de novo* mutations within each population.

First, we identified *de novo* mutations targeting the same genes in both monocultures and co-cultures. One such mutation was a loss-of-function mutation in the *gacS* gene that rapidly fixed or rose to high frequency (>75%) between generations 50 and 100 in both monocultures and co-cultures (Fig. 2, Fig. S2, Table S1-S2). Given the pervasive nature of this mutation and its rapid establishment in the population, it likely represents an adaptation to the culture conditions.

**Figure 2.**
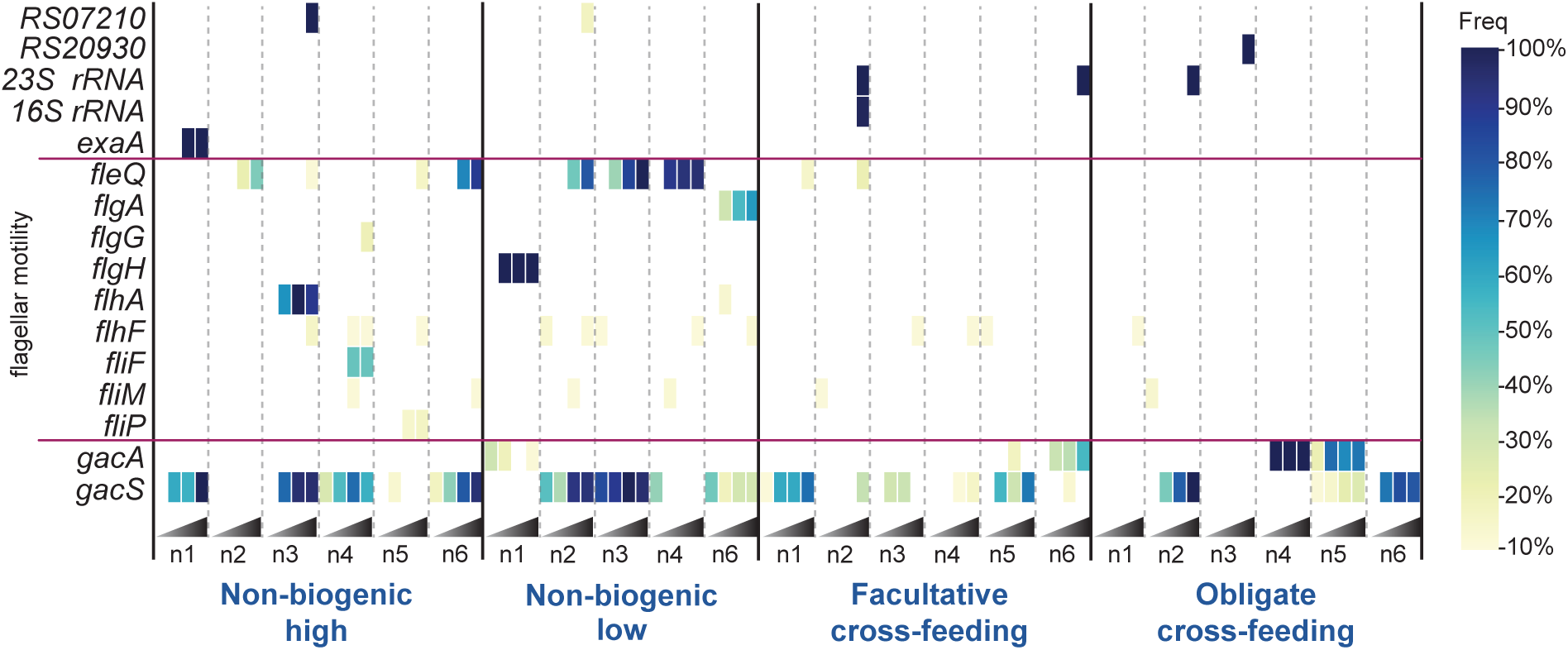
Unidirectional cross-feeding limits the acquisition of *de novo* mutations in flagellar genes in *P. putida.* Heatmap representing the frequency of *de novo* mutations in the evolved population of *P. putida* for all evolved lineages (n1 to n6) in each condition at generations 50, 100, 150, and 200 (represented from left to right for all lineages). Genes with a frequency above 10% have been plotted. Note that replicate n5 in the non-biogenic low condition went extinct during the experiment.

We also identified mutations that were exclusive to either monocultures or co-cultures. Notably, mutations targeting the flagellar gene cluster were observed at high frequency (>75%) in both (high and low non-biogenic) monoculture conditions (Fig. 2 and Fig. S2). These mutations were associated with loss-of-function changes, such as nonsense mutations and indels (Table S1-S2). Notably, mutations in the flagellar operon reached high frequency early in the evolution experiment (generations 50-150), indicating that rapid selective sweeps occurred when *P. putida* relied on a non-biogenic carbon source. Although flagellar mutations also occurred in co-culture conditions (four of six populations in facultative cross-feeding and one of six populations in obligate cross-feeding), they were present at a much lower frequency (22% on average) than in monocultures. Furthermore, they were detected only at generation 200 (Fig. 2 and Fig. S2), indicating their emergence was delayed relative to monocultures. Notably, flagellar mutations were less abundant and occurred at lower frequencies in obligate cross-feeding conditions than in facultative cross-feeding conditions (Tables S1 and S2).

These results highlight that the interaction strength determines the extent to which the flagellar operon is under selective constraint.

We then identified high-frequency mutations (either fixed in the population or present at frequencies above 50%) specific to the cross-feeding conditions. These mutations targeted the following genes: a gene encoding a transposase (*RS20930*), two RNA genes (23S rRNA and 16S rRNA), and the *gacA* gene encoding a UvrY/SirA/GacA family response regulator transcription factor (Fig. 2, Fig. S2, Tables S1-S2).

Overall, loss-of-function mutations in flagellar genes rapidly fixed in *P. putida* populations evolving on a non-biogenic carbon source but were selected against under facultative or obligate cross-feeding. These findings suggest that flagellar motility (or other associated traits) is essential for growth on biogenic carbon sources.

### Enhanced motility evolves under cross-feeding conditions

Based on the underrepresentation of loss-of-function mutations in flagellar genes under cross-feeding conditions (Fig. 2), we hypothesized that flagellar motility is important for cross-feeding. For example, flagellar motility may enable *P. putida* to swim toward the transient bursts of benzoate released by nonmotile *A. johnsonii*. To explore this possibility, we used a high-throughput fluorescence microscopy assay to assess *P. putida*’s swimming behavior in the presence of *A. johnsonii* (Fig. 3A). An ancestral *A. johnsonii* population was mixed at a 1:1 ratio with either the ancestral *P. putida* population or evolved *P. putida* populations from generation 200. Benzyl alcohol was supplied as the sole carbon source to model the obligate cross-feeding condition.

**Figure 3.**
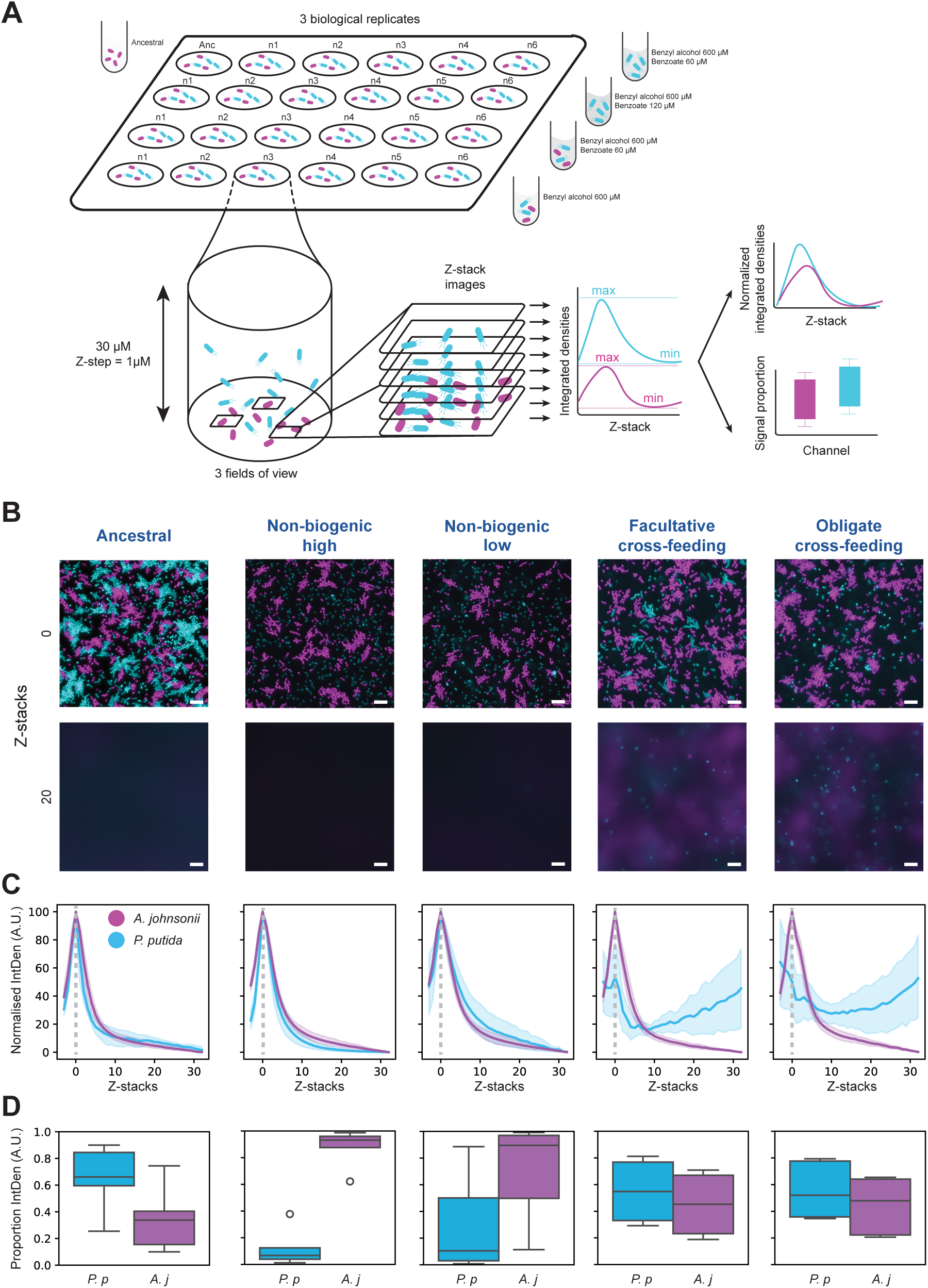
Unidirectional cross-feeding interactions promote the acquisition of enhanced motility in *P. putida.* (*A*) Design of the high-throughput microscopy assay used to visualize and quantify *P. putida’*s motility in the presence of *A. johnsonii*. (*B*) Representative images of the co-culture microscopic assay for ancestral and evolved lineages of *P. putida* (cyan) in the presence of ancestral *A. johnsonii* (magenta). Top row correspond to the images from to the bottom of the well (at the glass surface), and the bottom row correspond to images at z plane at 20 μm above the bottom of the well. *A. johnsonii* expressing RFP is shown in magenta, and *P. putida* expressing GFP is shown in cyan. Scale bar represents 10 μm. (*C*) Representative example of normalized integrated density of fluorescent signal corresponding to *A. johnsonii* (magenta), and *P. putida* (cyan) for all conditions tested across the z range (from 0 to 30 μm above the bottom of the well with an image taken with a 1 μm increment). Dark lines represent the median values from 3 different fields of view per lineage tested, and light areas represent the 95% CI. (*D*) Representative example of the proportion of integrated density fluorescent signal across the z-range between *P. putida* (cyan) and *A. johnsonii* (magenta) for all conditions and all lineages.

When ancestral populations of *A. johnsonii* and *P. putida* were mixed, most signals from both cell types were observed near the bottom of the well, and swimming was negligible across the z range (Fig. 3A-B and Fig. S3A, Movie S1). This observation confirms that under static co-culture conditions, only a small fraction of ancestral *P. putida* cells exhibit swimming behavior, while the majority remain associated with non-motile *A. johnsonii* at the bottom of the well. Similarly, when *P. putida* evolved lineages from the monoculture conditions were combined with the ancestral *A. johnsonii* population, most of the signal was concentrated at the bottom of the wells (Fig. 3A-B and Fig. S3A. Movies S2-3). This pattern is consistent with the loss-of-function mutations in the flagellar operon in these lineages (Fig. 2). However, the proportion of monoculture-evolved *P. putida* relative to *A. johnsonii* was significantly lower than in the ancestral population, indicating a potential fitness trade-off in the presence of *A. johnsonii* (Fig. 3C and Fig. S3B). In contrast, when co-culture evolved *P. putida* populations (facultative and obligate cross-feeding) were mixed with the ancestral *A. johnsonii* population, *P. putida* cells were detected across the z range (cyan dots at Z-stack 20 in Fig. 3A-B and Fig. S3A, Movies S4-5), indicating that a large fraction of cells exhibited swimming behavior. Time-lapse imaging further confirmed that *P. putida* populations evolved under cross-feeding were dominated by actively swimming individuals (Movie S6), whereas fewer cells were actively swimming in the ancestral population (Movie S5). These findings indicate that populations evolved under cross-feeding exhibit enhanced motility, characterized by a larger fraction of actively swimming cells. Furthermore, under these cross-feeding conditions, *P. putida* coexisted with *A. johnsonii* at a proportion similar to that of the ancestral population, indicating that the interaction remained stable throughout the experiment (Fig. 3C and Fig. S3B). Overall, these results suggest that selection under cross-feeding conditions not only maintains functional motility but may also enhance it.

### The fitness effect associated with specific mutations is context-dependent

Based on the genotypic characterization of evolved lineages (Fig. 2) and swimming behaviors (Fig. 3) we hypothesize that loss-of-function mutations in flagellar genes (enriched in monocultures) confer a fitness advantage in monoculture but a fitness disadvantage in co-culture with *A. johnsonii*. Conversely, we hypothesize that mutations promoting foraging-associated traits (e.g., motility and chemotaxis), as well as other mutations enriched under cross-feeding conditions, confer a fitness advantage in co-culture.

To test these hypotheses, we isolated clones with single mutations enriched in either monoculture or co-cultures (Table S1). In addition, we isolated clones displaying enhanced motility (mSwim). We confirmed that these mutants retained the phenotypic characteristics previously observed in populations from which they were isolated.

Specifically, the *fleQ* and *flgA* mutants exhibited no motility (Fig. S4) and higher yields (Fig. S5A) and, whereas the mSwim mutant exhibited enhanced motility (Fig. S4 and S5B). In parallel, we genetically engineered a *P. putida* mutant with a clean deletion of the *cheA* gene, which encodes the chemotaxis regulator (ΔcheA), to test whether directed motility could provide a fitness advantage to *P. putida* under cross-feeding conditions.

We quantified the fitness effects of these mutations in monoculture and co-culture conditions using head-to-head competition experiments between mutants and the *P. putida* ancestor (Fig. 4A). The *fleQ* and *flgA* mutants had a large fitness advantage in monoculture and outcompeted the ancestor within 24h, regardless of their initial ratio (Fig. 4B and Fig. S5C). This high fitness advantage may explain why loss-of-function mutations rapidly swept to fixation under monoculture conditions during our evolution experiment. In contrast, these *fleQ* and *flgA* mutations conferred no fitness advantage in co-culture with *A. johnsonii* and even imposed a fitness cost depending on their initial frequency (Fig. 4C and Fig. S5D). This suggests that the loss of flagellar motility affects *P. putida*’s competitive ability when relying on cross-feeding interactions.

**Figure 4.**
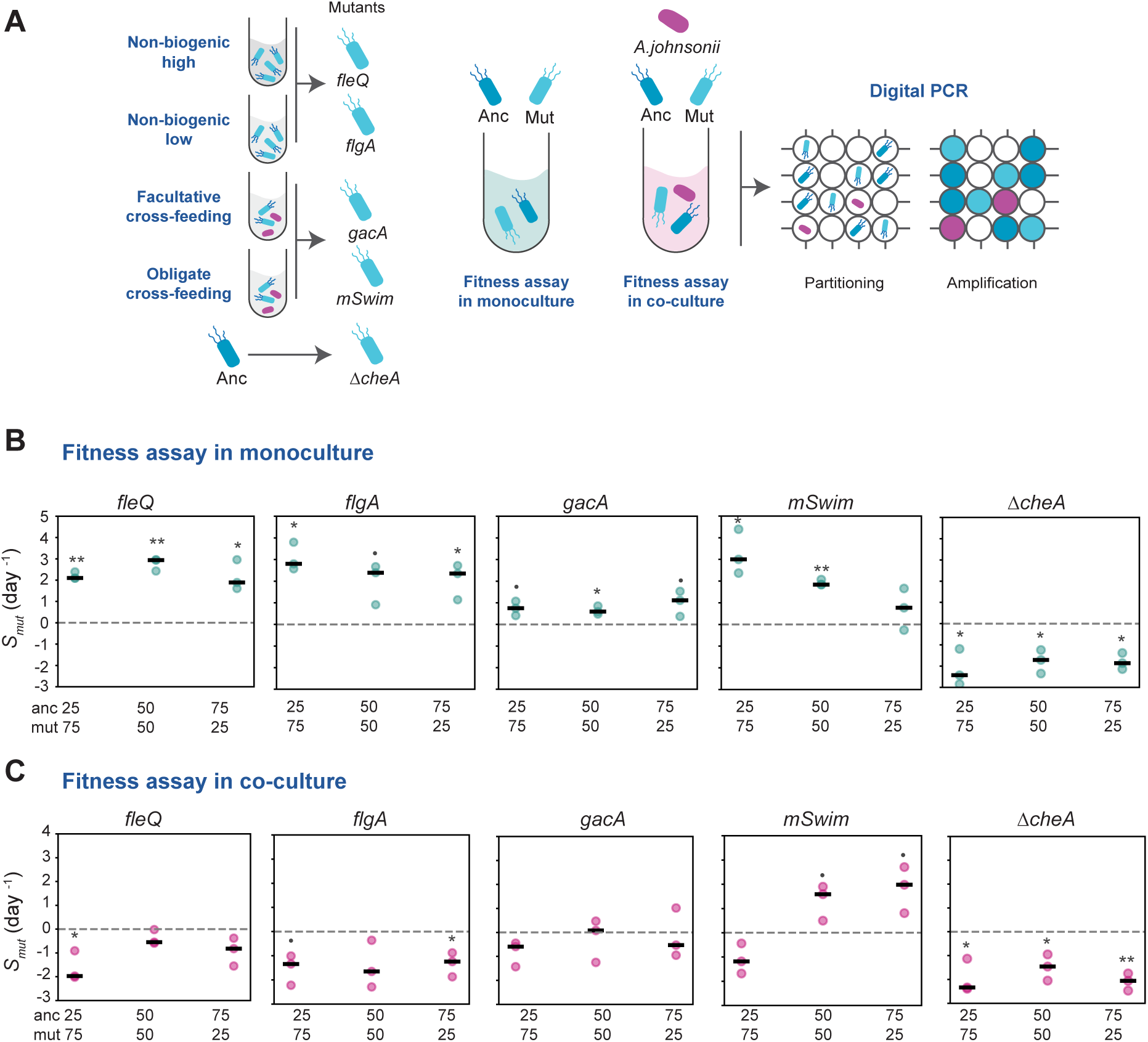
The fitness effect associated with specific *de novo* mutations is context dependent. (*A*) A library of mutants (left panel) was generated by isolating clones with mutations representative of monoculture conditions (*fleQ* and *flgA* mutants) and cross-feeding conditions (*gacA* and mSwim mutants), and by knocking down the *cheA* gene (Δ*cheA*). Relative fitness assays were performed using digital PCR (right panel). (*B* and *C*) Scatterplot representing the selection coefficient of mutant at different initial starting ratios in monoculture (panel *B*) and in co-culture with *A. johnsonii* (panel *C*). The black line represents the median value, and dots represent three biological replicates. Asterisks denote significance from a one-sample t-test testing whether the selection coefficient differs from zero (** *P* <0.01, * *P* <0.05). The dot (•) indicates marginal significance (0.05 < *P* < 0.1).

The mutations representative of cross-feeding conditions (*gacA,* mSwim) showed a more nuanced trend. The *gacA* mutation conferred a slight fitness advantage under monoculture conditions (Fig. 4B and Fig. S5C). Under co-culture conditions, the *gacA* mutation had no significant fitness advantage (Fig. 4C and Fig. S5D). Therefore, the fitness effect of this mutation cannot fully explain why it was observed at high frequency exclusively in cross-feeding conditions (Fig. 2). In turn, the mutant associated with a gain of motility (mSwim) had a fitness advantage in both monoculture and co-culture conditions but in a frequency-dependent manner (Fig. 4B-C and Fig. S5C-D). In monoculture, the mSwim population had a significant fitness advantage when starting from higher and equal proportions with the ancestor (75_mut_:25_anc_ and 50_mut_:50_anc_; Fig. 4B and Fig. S5C). This effect was reversed in co-culture conditions, in which the mSwim population showed a marginal fitness advantage when starting from lower or from equal proportions (Fig. 4C and Fig. S5D). Finally, the ΔcheA mutant showed a significant fitness disadvantage under both monoculture and co-culture conditions (Fig. 4B and Fig. S5C). This indicates that, directed motility provides a fitness advantage to *P. putida* in cross-feeding conditions as well as in well-shaken environments supplied with external resources.

## Discussion

Our study revealed two distinct evolutionary trajectories in the consumer species (*P. putida*), determined by the presence or absence of a producer species (*A. johnsonii*). In monoculture, adaptation to nutrient scarcity occurs through loss-of-function mutations in flagellar genes, accompanied by substantial growth improvements. In contrast, under both facultative and obligate unidirectional cross-feeding, adaptation involves maintenance and enhancement of flagellar motility.

Our first major finding is that selection favors the loss of flagellar motility when the consumer species evolves in isolation with externally supplied non-biogenic resources (Fig. 2 and 3). Several experimental evolution studies have shown that loss-of-function mutations in the flagellar operon rapidly arise in well-mixed environments (12–14). A common explanation for this phenomenon is that flagellar motility is not needed in well-mixed environments supplemented with external resources. Given that motility is highly costly (15), reducing or losing motility saves energy that can be reallocated for other cellular functions, such as biomass production (16). In our study, loss-of-function mutations occurred in genes involved in both the regulation of flagellar motility (e.g., *fleQ* encoding the master regulator FleQ) and flagellar assembly and structure (e.g., *flgA* and *flgH* encoding proteins involved in the formation of the P- and L-rings of the flagellar basal body). We demonstrated that at least two of these flagellar mutations (one regulatory and one structural) were highly advantageous under monoculture conditions. These large fitness effects are perhaps unsurprising given that benzoate concentrations were low in both resource treatments (non-biogenic ’low’ and ’high’). Under such strong resource limitation, selection is expected to favor suppression of energetically costly motility. Unexpectedly, however, cells with enhanced motility (mSwim) also exhibited a fitness advantage under monoculture conditions, seemingly contradicting our hypothesis that motility is costly under resource limitation. These observations should be interpreted with caution. Given that the genetic basis of the enhanced-motility phenotype remains unresolved, we used a mutation (*gacS*) to distinguish evolved mSwim clones from the ancestor. However, *gacS* was observed at high frequency in both monoculture and co-culture conditions, suggesting that it may represent an adaptation to these culture conditions. Thus, these putatively beneficial mutations may act as a confounding factor underlying the observed effects.

Our second major finding is that selection favors the maintenance of functional motility when consumers evolve under cross-feeding conditions. This conclusion is supported by experiments showing that nonmotile clones (i.e., *fleQ* and *flgA* mutants) exhibited a fitness disadvantage in competition with the motile ancestor in the presence of the producer species. Thus, the fitness benefits conferred by these mutations in monoculture were lost under obligate cross-feeding conditions, where they are neutral or deleterious. This pattern is consistent with antagonistic pleiotropy, in which the fitness effects of mutations depend on ecological context, and may help explain why loss-of-function mutations in flagellar genes were frequently enriched in monoculture but occurred at much lower frequency under cross-feeding conditions. In addition to retaining motility, lineages evolved under cross-feeding conditions displayed enhanced motility relative to the ancestor. Why would enhance motility be selected under cross-feeding conditions? We reason that by releasing benzoate in pulses, the producer creates transient resource gradients. By retaining/enhancing motility, the consumer could more efficiently swim toward the producer species and consume high-energy resources excreted by the producer. Such dynamics have been observed in systems with transient resource patches, where selection favors foraging-associated traits such as chemotaxis, swimming, and surface attachment (17–19). One consideration is whether resource gradients persist under our shaking conditions (150 rotations per minute, rpm). We consider this is plausible, as we have observed that producer cells do form aggregates under 150 pm shaking conditions, potentially generating resource gradients. Even in the absence of strong gradients, Scarinci and Sourjik (2023), have shown that flagellar motility provides an advantage in relatively fast shaking conditions (300 rpm) when a cross-feeding species is present (20). They hypothesize this is because motility increases encounter rates between species and promotes cell-cell adhesion (20).

Taken together, our results indicate that resources origin (biogenic and non-biogenic) is an important factor driving the divergent evolutionary trajectories observed in consumer populations. One potential limitation is that it remains difficult to disentangle the extent to which the observed differences in consumer evolution arise solely from resource heterogeneity due to cross-feeding versus additional biotic effects exerted by the producer on the consumer. Moreover, although it would have been ideal to harvest benzoate excreted by the producer for use in monoculture treatments to ensure that the benzoate is chemically identical, this is challenging in practice. For example, supernatant-based approaches are complicated by the fact that benzoate is an intermediate by-product that is subsequently reutilized and depleted by the producer, making accurate timing of peak excretion difficult. Future studies could begin disentangling the relative contribution of biotic and abiotic factors by evolving consumer populations under pulsed resource availability in the absence of the producer species.

In conclusion, our results demonstrate that unidirectional cross-feeding can play a key role in shaping the evolutionary trajectories of the consumer. Loss of flagellar motility is selected in presence of externally supplied resources, whereas motility is maintained under cross-feeding conditions. This pattern persists even when facultative cross-feeding populations are exposed to non-biogenic carbon sources, a condition commonly encountered in natural environments, suggesting that biotic interactions strongly influence evolutionary outcomes. Overall, our findings highlight the importance of studying evolution within a community context.

## Materials and Methods

### Strains, plasmids, and primers

We used a well-characterized consortium of *P. putida* strain KT2440 and *A. johnsonii* strain C6 originally isolated from a creosote polluted aquifer in Denmark (8, 9, 21). The *P. putida* KT2440 strain used in this study is unable to metabolize benzyl alcohol (absence of TOL/pWW0 plasmid), has flagellar motility, and is resistant to gentamicin (gfp-Gm^r^ chromosomal cassette; 8). *A. johnsonii* strain C6 lacks flagellar motility and is resistant to streptomycin (22). All other strains, plasmids, and primers used in this study are listed in Table S3.

### Media & solutions

Strains were grown in FAB minimal medium containing (NH4)_2_ SO_4_ 15 mM, Na_2_HPO_4_ 33.3 mM, KH_2_PO_4_ 22 mM, NaCl 50 mM, CaCl_2_ 0.1 mM, MgCl_2_ 1 mM, and Fe-EDTA 0.01 mM, and supplemented with either benzyl alcohol or benzoate or both carbon sources. To minimize the presence of assimilable organic carbon (AOC), bacterial cultures and solutions were prepared and stored in muffled glassware, as described in Rodriguez-Verdugo et al. 2019 (9).

### Evolution experiment

We evolved *P. putida* either alone (in monoculture) or in co-culture with *A. johnsonii* for approximately 200 generations (30 days) under one of four experimental treatments:

1. Obligate cross-feeding: Co-cultures were supplemented with benzyl alcohol (600 μM) only. In this condition, benzoate (∼60 μM based on HPLC measurements; ref. 9) is produced by *A. johnsonii*.
2. Facultative cross-feeding: Co-cultures were supplemented with benzyl alcohol (600 μM) and benzoate (60 μM). Benzoate is both produced by *A. johnsonii* and supplied externally, yielding ∼120 μM total benzoate.
3. Non-biogenic low: Monocultures of *P. putida* were supplemented with benzyl alcohol (600 μM) and benzoate (60 μM).
4. Non-biogenic high: Monocultures of *P. putida* were supplemented with benzyl alcohol (600 μM) and benzoate (120 μM).

To start the evolution experiment, *P. putida* and *A. johnsonii* were revived by streaking glycerol stocks onto LB agar plates supplemented with gentamicin (10 µg/mL) and streptomycin (64 µg/mL), respectively, and incubated at 30°C. A single colony from each species was randomly selected and inoculated into 10 mL of FAB medium + benzyl alcohol (600 μM) for *A. johnsonii* and benzoate (60 μM or 120 μM) for *P. putida*. Cultures were incubated at 26°C with shaking at 150 rpm for 24 hours. Monocultures were launched by inoculating 100 µL of acclimated cultures into 9.9 mL of FAB medium with the specified carbon sources. Co-cultures were initiated by mixing 50 µL from each species culture into 9.9 mL of FAB medium. To ensure equal starting ratios, all co-cultures were plated immediately after inoculation, and colony counts confirmed a 1:1 ratio of *P. putida* to *A. johnsonii*. Glycerol stocks (20%) of these ancestors were stored at -80°C.

We conducted the evolution experiment using six independent replicate populations (lineages) per treatment, propagated under identical conditions. Cultures were transferred daily by inoculating 0.1 mL of each population into 9.9 mL of fresh FAB minimal medium supplemented with the appropriate carbon sources. Cultures were incubated at 26°C with shaking at 150 rpm. To prevent cross-contamination during daily transfers, pipettes were cleaned with ethanol between each transfer. In addition, populations were plated weekly to monitor potential contamination.

To track population dynamics during the experiment, all replicates were plated onto selective LB agar plates supplemented with gentamicin or streptomycin at generations 0, 50, 100, 150, and 200. Glycerol stocks of these time points were stored at -80°C for subsequent phenotypic and genomic analyses.

### Quantification of growth-associated traits

To compare growth-associated traits between the ancestral and evolved populations of *P. putida*, we used a previously described 96-well plate growth curve assay (11). Briefly, evolved populations were revived by resuspending 100 μL of glycerol stock in 10 mL of FAB medium containing the same carbon source used during the evolution experiment. Cultures were incubated overnight at 26 °C with shaking at 150 rpm and then plated onto LB agar supplemented with the appropriate antibiotics. Plates were incubated overnight at 30 °C, and 60 colonies were randomly selected and resuspended in 200 μL of 1% MgSO4 in a 96-well plate. Two microliters of each resuspension were inoculated into 198 μL of FAB medium supplemented with 600 μM benzoate. The 96-well plates were sealed with high-vacuum grease to prevent evaporation, and optical density at 600nm was measured every 5 minutes for 24 h at 30°C with constant shaking using a BioTek plate reader. Growth parameters (total yield of growth, maximum growth rate, half-saturation constant, and maximum uptake rate) were estimated from OD measurements using a Python script adapted from (9). For each growth parameter, the median value across 60 colonies was calculated and divided by the median of 60 ancestral colonies to quantify growth relative to the ancestor.

### *De novo* mutations characterization

To characterize *de novo* mutations in the evolved population of *P. putida,* we performed short read sequencing of evolved populations. Briefly, a glycerol stocks were thawed and 100 μL was inoculated in 10 mL of FAB medium containing the same carbon source used in the evolution experiment.

Cultures were incubated overnight at 26°C with shaking at 150 rpm. Cells were spin down by centrifuging 5 mL of culture for 5 min at 13000 rpm. Genomic DNA was extracted using the Promega Wizard® HMW DNA Extraction Kit. Libraries were prepared using Nextera primers with a low-reaction-volume protocol adapted from Illumina (23). Sequencing was performed on an Illumina NovaSeq 6000 (S4 flow cell) using 150-bp paired-end reads with 5% PhiX Spike-in.

Read quality was assessed using FastQC (v0.11.9) and MultiQC (v1.0) (24), and adapter sequences were removed using TrimGalore (v0.6.7) (25). Mutations were identified by aligning reads to the reference genome of *P. putida* KT2440 (reference NC_002947) using Breseq (v0.36.0) (26). We used polymorphism mode to call variants. Each sample was sequence to a depth of ∼100x coverage, allowing detection of mutations at frequencies above 10% in the population. *De novo* mutations were defined as those uniquely present in evolved populations and absent from the ancestor.

### Imaging and quantification of motility at the population level

We conducted a high-throughput fluorescence microscopy assay to characterize swimming behaviors in *P. putida* (Fig. 3A). To prepare the starter cultures for the microscopy assays, glycerol stocks of ancestral or evolved populations were thawed and 100 μL was inoculated into 10 mL of FAB medium supplemented with the same carbon source used during the evolution experiment. Cultures were incubated overnight at 26°C with shaking at 150 rpm. Cells were harvested by centrifugation for 10 min at 4,000 rpm and resuspended in 1 mL of FAB medium supplemented with 1mM benzyl alcohol. Optical density at 600 nm was measured and normalized to 0.1.

To visualize *A. johnsonii* (RFP) and *P. putida* (GFP) in co-culture, we used a glass-bottom 24-well plate with #1.5 cover glass (Cellvis P24-1.5H-N). Wells were pre-filled with 1960 mL of FAB medium supplemented with 1 mM benzyl alcohol. To obtain a starting OD of 0.001, 20 μL of normalized starter cultures of *A. johnsonii* and *P. putida* were added. Plates were incubated in a humidified chamber at 26 °C and imaged using a Leica THUNDER 3D epifluorescence microscope (inverted) fitted with LED8 light source and a K8 (4.2 MP) sCMOS camera. GFP and RFP fluorescence over a Z range of 35 μm was acquired at 24, 48, and 72 h post-inoculation with a 40X/0.95 NA air objective (Voxel size: 0.16 x 0.16 x 1 µm). Time-lapse images were additionally collected at Z positions corresponding to Z = 0 μm (the bottom of the chamber, at glass-medium interface), Z = 10 μm (mid-depth), and Z = 30 μm (deep in the chamber, above the glass surface).

Images were processed and analyzed using Leica LASX, FIJI, and Python using the Scikit-Image plugin (27). Quantification of the integrated density over the Z range, calculation of the maximum projection intensity, and determination of the overall proportion of each signal for both GFP and RFP were performed using Python.

### Isolation and selection of clonal populations with specific *de novo* mutations

We isolated clones carrying mutations specific to either monoculture (*fleQ* and *flgA)* and co-culture (*gacA*) conditions (Table S2), as well as clones exhibiting enhanced motility (mSwim) from evolved populations. These clones were isolated from 200-generations evolved lineages harboring these mutations at high frequency (>50%): non-biogenic low (replicate 4) for *fleQ* ; non-biogenic low (replicate 6) for *flgA*; and obligate cross-feeding (replicate 4) for *gacA*. In the case of clones with enhanced motility (mSwim), we were unable to identify the genetic basis of this phenotype, therefore, we isolated clones from the obligate cross-feeding population (replicate 6) that contained a mutation in the *gacS* gene. This mutation was not specific to cross-feeding conditions (occurring in both monoculture and co-culture) but enabled the design of probes to distinguish evolved clones from the ancestral *P. putida* population.

Populations were revived by inoculating 100 µl of the glycerol stock into FAB medium supplemented with 600 μM benzoate. After 24h of incubation at 30°C at 150 rpm, cultures were diluted and plated onto LB agar supplemented with gentamycin (10 μg/mL). Sixteen colonies were isolated and screened by PCR targeting the gene of interest, and amplicons were sequenced to confirm the presence of the expected *de novo* mutations (Table S1-2). Confirmed clones (called mutants) were then assayed for growth traits (as described above) to verify that they recapitulated previously observed phenotypes. In addition, we performed swimming assays of these clones and the ancestor using 0.3% (wt/vol) soft agar as in (11).

### Construction of the *ΔcheA* mutant

To generate a clean deletion of the *cheA* gene in the ancestral *P. putida* background, we used the Gibson assembly technology.

Cassettes corresponding to the kanamycin resistance gene (*kan)* from the pDK4 plasmid and regions upstream (500 bp upstream of the *cheA* start codon) and downstream (500 bp downstream of the *cheA* stop codon), were amplified with flanking regions to anneal to pTW475 (SmaI site). Gibson assembly was then performed, and colonies were screened by sequencing to confirm the insertion of these cassettes into pTW475. A mating protocol between the *E. coli* strain carrying this plasmid and the ancestral strain of *P. putida* was performed. Recombinant *P. putida* clones were screened, and PCR products were sequenced to confirm the deletion of the *cheA* gene by the insertion of the *kan* gene.

### Relative fitness assays using digital PCR

To quantify the fitness effects of specific *de novo* mutations in different ecological contexts, we conducted ’head-to-head’ competition experiments between mutants and ancestor of *P. putida* in different conditions (Fig 4A).

First, we revived the mutants and *P. putida* and *A. johnsonii*’s ancestors of by inoculating 100µL glycerol stocks into FAB medium supplemented with 1 mM benzoate (for *P. putida*) and 1 mM benzyl alcohol (for *A. johnsonii*) and incubating them overnight at 26°C with shaking 150 rpm. Cultures were normalized to OD 1 to standardize the starting ratios. Competition assays in monoculture (mutant vs *P. putida* ancestor) were performed in 10 mL of FAB medium supplemented with 1 mM benzoate as the sole carbon source. Competition assay in co-culture (mutant vs *P. putida* ancestor in the presence *A. johnsonii* ancestor) were performed in 10 mL of FAB medium supplemented with 1 mM of benzyl alcohol as the sole carbon source. In both conditions, *P. putida* mutants and ancestor were mixed at 25:75, 50:50, and 75:25 starting ratios. For co-cultures, *A. johnsonii* and *P. putida* were mixed at a 50:50 starting ratio. Cultures were incubated for 24h at 30°C with constant shaking (150 rpm). The proportion of the mutant vs ancestor at the start and the end of the 24-hours experiment was quantified using digital PCR assays. Genomic DNA was extracted from pellets (obtained by centrifuging the cultures for 10 min at 4000 rpm) using the Promega Wizard® HMW DNA Extraction Kit. We use 2 ng of gDNA input for the digital PCR assays.

The digital PCR assay was designed using the QuantStudio Absolute Q Digital PCR system. First, a set of primers and probes was designed to amplify the genomic region surrounding the specific mutations and to anneal to either the ancestral or the mutated genomic region. Using QuantStudio software, the threshold was adjusted to remove the background signal, and positive wells were quantified for the targeted pair of probes. Based on the total number of wells quantified per digital PCR assay, the number of positive wells was normalized, and the proportion of ancestral vs mutant was quantified.

The selection coefficient for a given mutant (*s_mut_*), defined as the relative fitness difference between a mutant and the ancestor, was calculated using the following formula: [inlin e], where *p_0_* and *p_t_* are the mutant frequencies at the beginning and end of the competition experiment, and *q_0_* and *q_t_* are the ancestor frequencies at the beginning and end of the competition experiment. Competition experiments were performed in triplicate, and the mean selection coefficient was calculated. A selection coefficient significantly greater than 0 indicated the mutant had a selective advantage relative to the ancestor, whereas a selection coefficient less than 0 indicated that the mutant had a selective disadvantage relative to the ancestor.

### Data and statistical analyses

All experiments described above were repeated at least three times to obtain a minimum of three biological replicates (n=3; additional replicated are noted where applicable). Data were processed and analyzed with Python (v3.11.5) in JupyterLab with the Numpy (28), Pandas (29), and Bioscikit packages. Graphical visualizations were generated using Matplotlib (30) and Seaborn (31). Statistical analyses were performed with the Statmodels package. ANOVA and Tukey’s HSD tests were performed to compare multiple pairs of group means from our dataset.

## Data availability

Raw sequences, images from the microscopic assays, and data from the growth curve and fitness assays will be available upon request. The code used to analyze and interpret datasets from this study is available on GitHub (github.com/Pirneote).

## Acknowledgments

We thank Dr. Travis Wiles for kindly providing plasmids pTW475 and pKD4 used to generate the Δ*cheA* strain. We thank Dr. Jennifer Martiny, Dr. Travis Wiles, Dr. Alexander Chase, Dr. Shivashankar Othy, and members of the Rodriguez-Verdugo and Martiny lab for providing helpful comments on the research and manuscript.

This work was funded by a grant from the United States National Science Foundation (DEB-2234627) to ARV. This work utilized resources of the UCI Genomics Research and Technology Hub (GRT Hub) parts of which are supported by NIH grants to the Comprehensive Cancer Center (P30CA-062203) and the UCI Skin Biology Resource Based Center (P30AR075047) at the University of California, Irvine, as well as to the GRT Hub for instrumentation (1S10OD010794-01and 1S10OD021718-01).

