## Supplementary Materials for "Microbial cross-feeding interactions reshape evolutionary trajectories of consumers by preserving motility"

### Supplementary figures

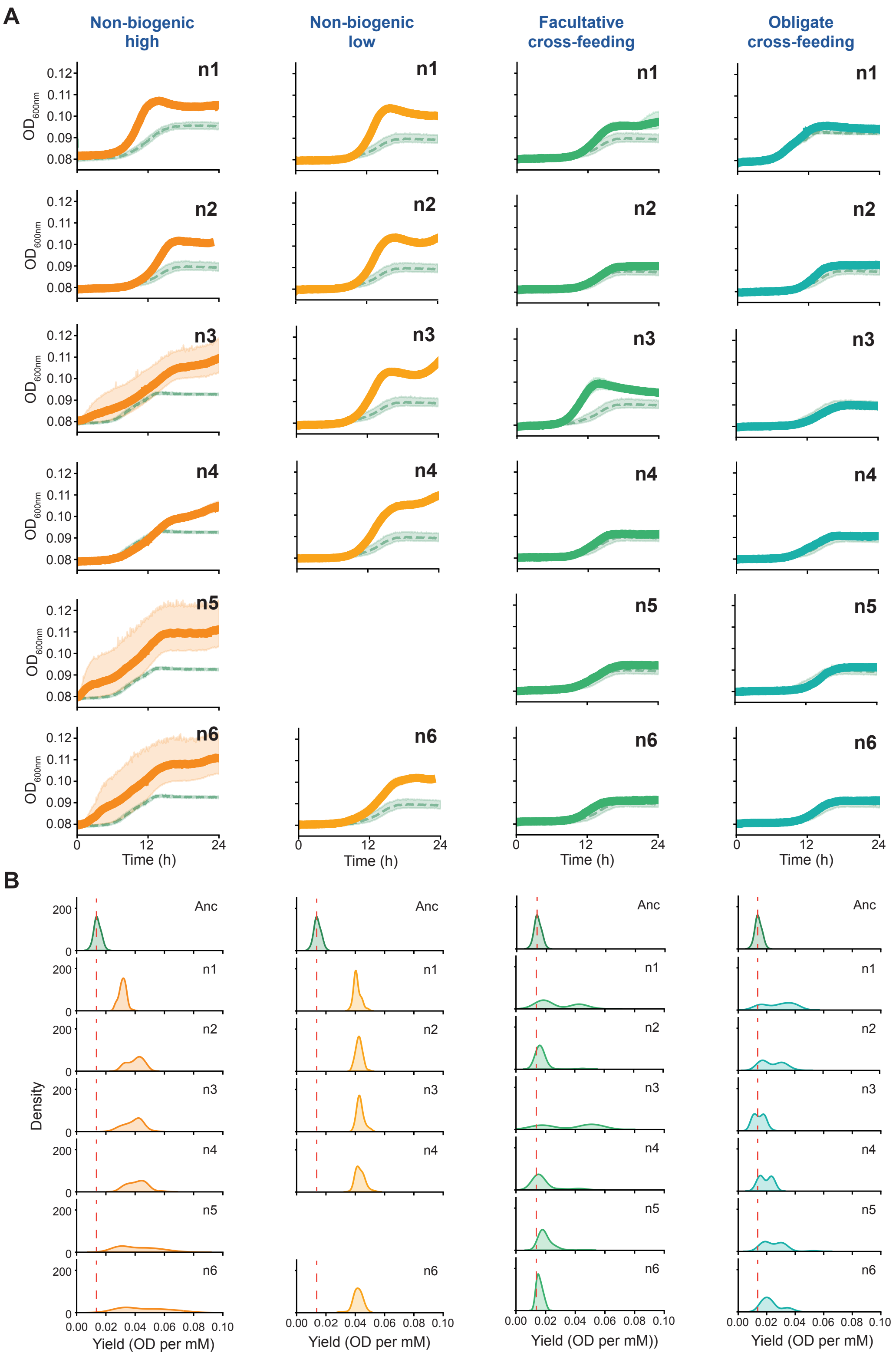

**Figure S1.** (*A*) Line plots representing the optical density ($OD_{600}$) over time of 60 colonies for the ancestral population and each replicate of the evolved populations at generation 200. Each line represents the median, and the shaded area denotes the 95% CI across 60 colonies sampled from the ancestral and evolved populations. The ancestral population is shown as a light green dashed line. Note that replicate n5 in the non-biogenic low condition went extinct during the experiment. (*B*) Density plots of the final yield (OD per mM) of 60 colonies for the ancestral and each replicate of the evolved populations at generation 200. The red-dashed line represents the median final yield for the ancestral population. Populations of *P. putida* evolved in non-biogenic benzoate evolved toward higher-yield distributions, whereas populations evolved under cross-feeding conditions retained distributions similar to that of the ancestor.

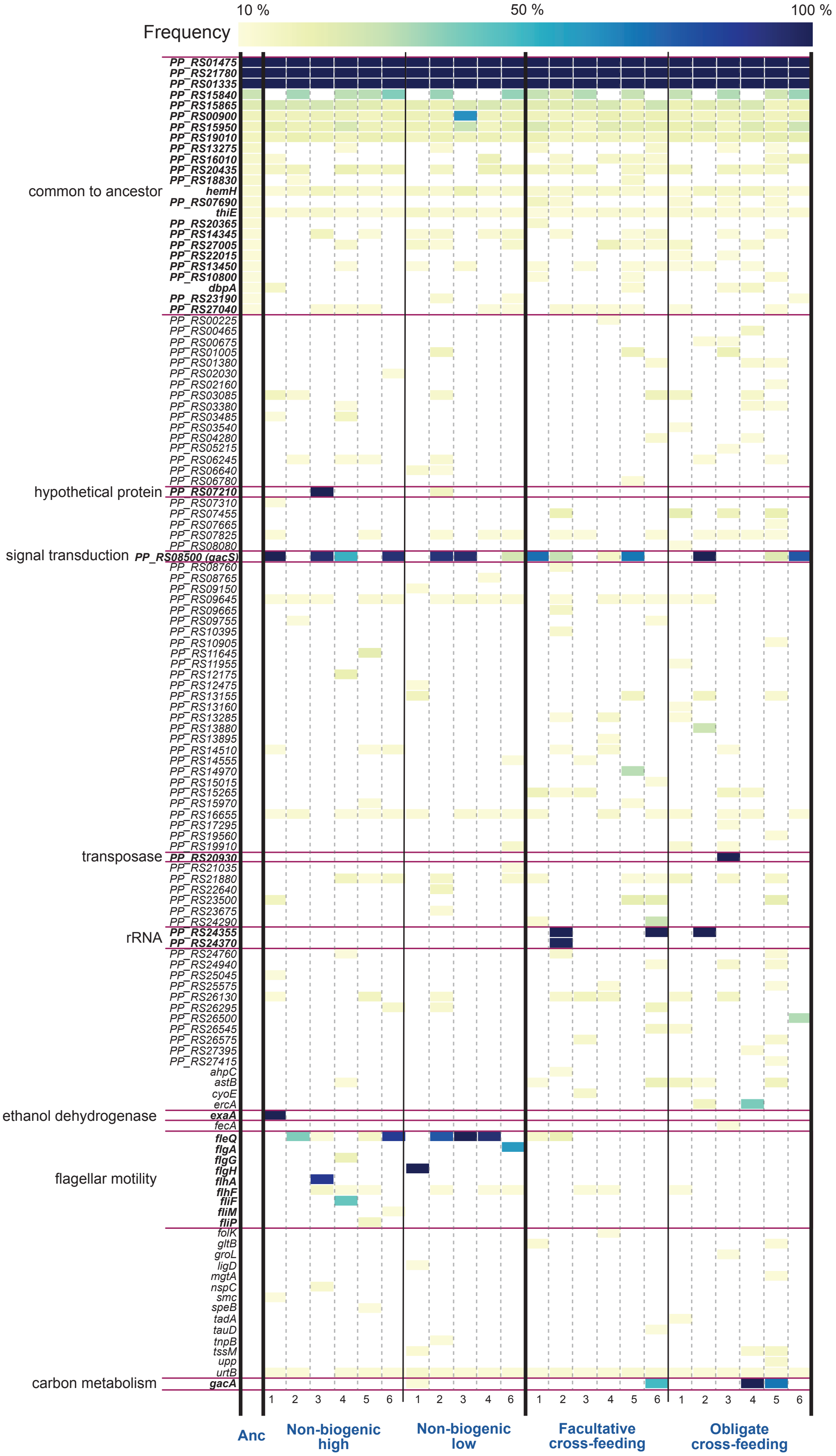
Frequency
10 %
50 %
100 %
common to ancestor
PP_RS01475
PP_RS21780
PP_RS01335
PP_RS15840
PP_RS15865
PP_RS00900
PP_RS15950
PP_RS19010
PP_RS13275
PP_RS16010
PP_RS20435
PP_RS18830
hemH
PP_RS07690
thiE
PP_RS20365
PP_RS14345
PP_RS27005
PP_RS22015
PP_RS13450
PP_RS10800
dbpA
PP_RS23190
PP_RS27040
PP_RS00225
PP_RS00465
PP_RS00675
PP_RS01005
PP_RS01380
PP_RS02030
PP_RS02160
PP_RS03085
PP_RS03380
PP_RS03485
PP_RS03540
PP_RS04280
PP_RS05215
PP_RS06245
PP_RS06640
PP_RS06780
hypothetical protein
PP_RS07210
PP_RS07310
PP_RS07455
PP_RS07665
PP_RS07825
PP_RS08080
signal transduction
PP_RS08500 (gacS)
PP_RS08760
PP_RS08765
PP_RS09150
PP_RS09645
PP_RS09665
PP_RS09755
PP_RS10395
PP_RS10905
PP_RS11645
PP_RS11955
PP_RS12175
PP_RS12475
PP_RS13155
PP_RS13160
PP_RS13285
PP_RS13880
PP_RS13895
PP_RS14510
PP_RS14555
PP_RS14970
PP_RS15015
PP_RS15265
PP_RS15970
PP_RS16655
PP_RS17295
PP_RS19560
PP_RS19910
transposase
PP_RS20930
PP_RS21035
PP_RS21880
PP_RS22640
PP_RS23500
PP_RS23675
PP_RS24290
rRNA
PP_RS24355
PP_RS24370
PP_RS24760
PP_RS24940
PP_RS25045
PP_RS25575
PP_RS26130
PP_RS26295
PP_RS26500
PP_RS26545
PP_RS26575
PP_RS27395
PP_RS27415
ahpC
astB
cyoE
ercA
ethanol dehydrogenase
exaA
fecA
flagellar motility
fleQ
flgA
flgG
flgH
flhA
flhF
fliF
fliM
fliP
folK
gltB
groL
ligD
mgtA
nspC
smc
speB
tadA
tauD
tnpB
tssM
upp
urtB
carbon metabolism
gacA
1 2 3 4 5 6 1 2 3 4 5 6 1 2 3 4 5 6 1 2 3 4 5 6
Anc
Non-biogenic high
Non-biogenic low
Facultative cross-feeding
Obligate cross-feeding

**Figure S2.** Heatmap showing mutation frequencies in evolved *P. putida* populations across all lineages under each condition at generation 200. Only genes with mutation frequencies above 10% are shown. *De novo* mutations were defined as those observed exclusively in evolved populations and absent in the ancestral population (i.e., mutations common to the ancestor were excluded from further analyses).

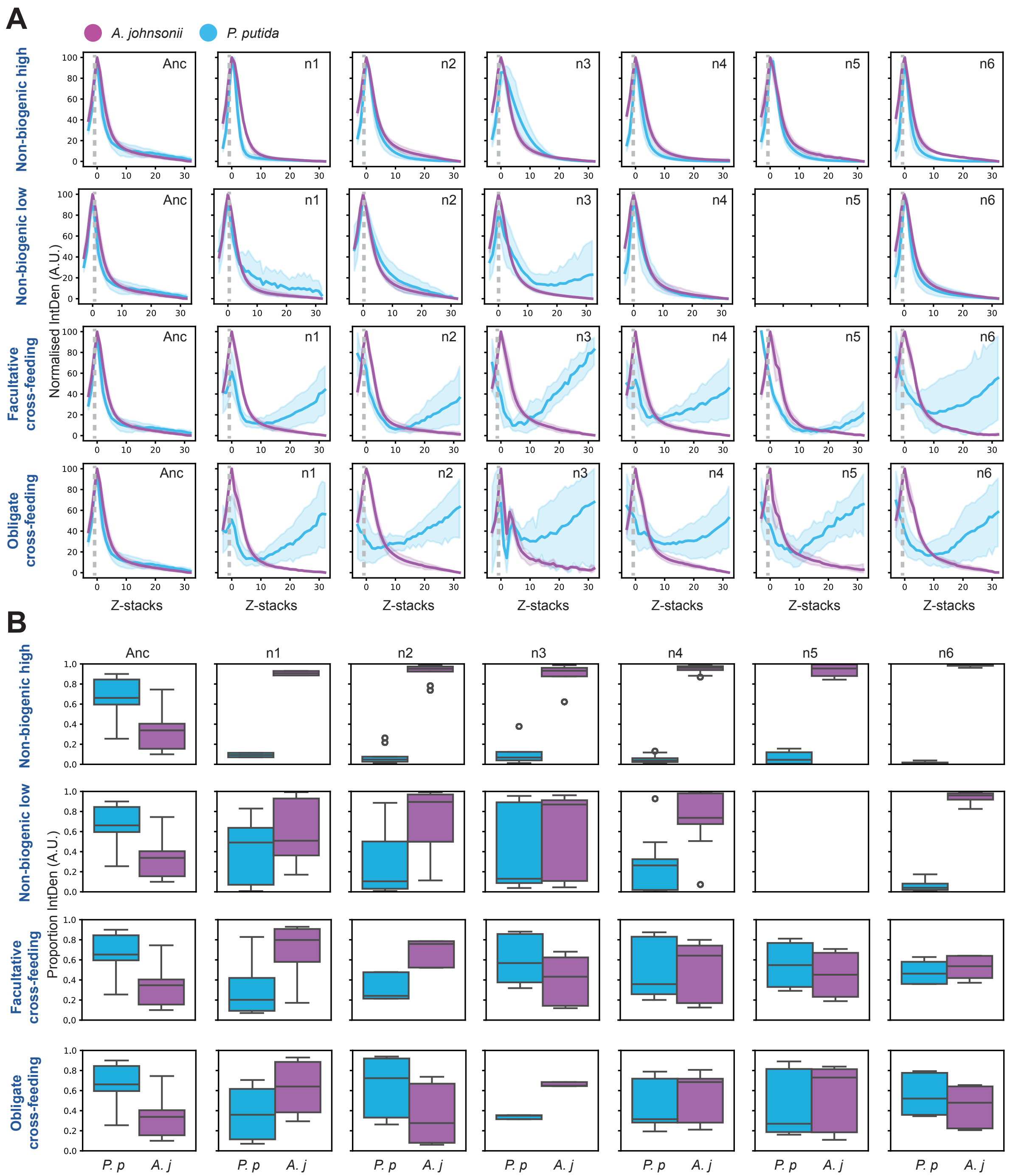


**Figure S3.** (*A*) Normalized integrated density fluorescent signal corresponding to *P. putida* (cyan), and *A. johnsonii* (magenta) for all conditions and all lineages tested across the z range (from 0 to 30 μm above the bottom of the well with images taken with 1 μm increments). Dark-colored lines correspond to the median values of 3 different fields of view per lineage tested, and light-colored areas correspond to the 95% CI. (*B*) Proportion of integrated density fluorescent signal across the z-range between *P. putida* (cyan) and *A. johnsonii* (magenta) for all conditions and all lineages.

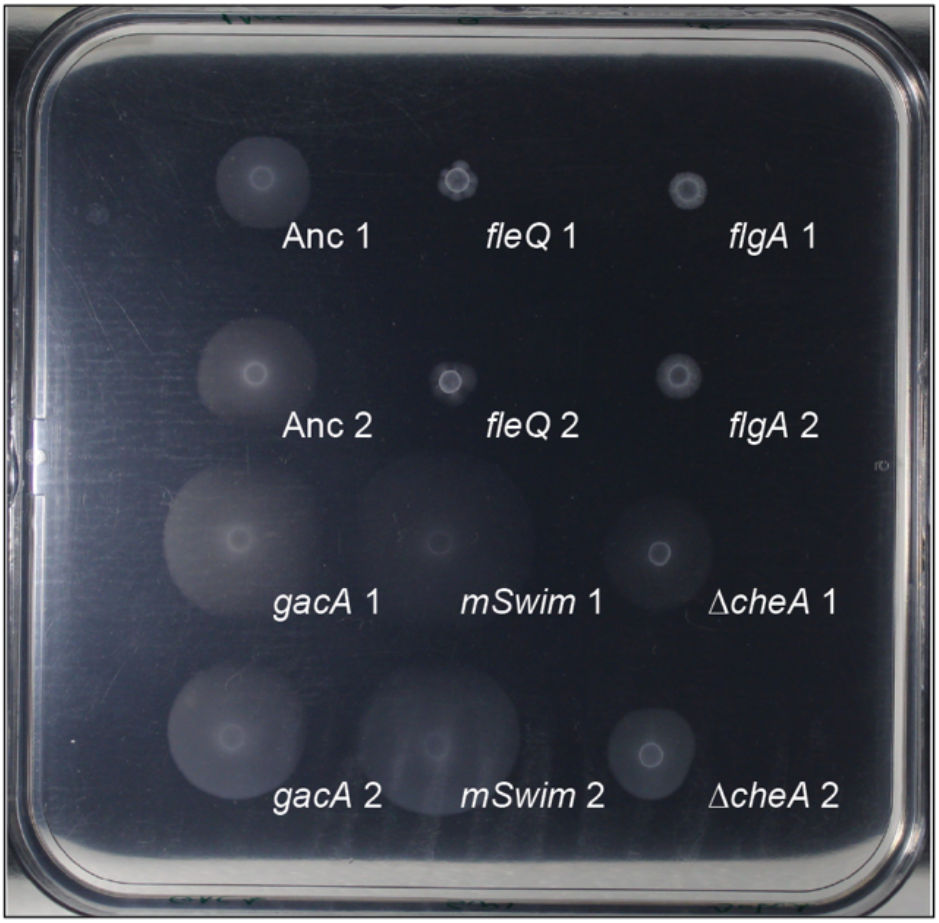


**Figure S4.** Swimming assay of clones with specific mutations or phenotypes isolated from evolved *P. putida* populations at 200 generations. Two technical replicates are shown, and three biological replicates were performed. Assays were conducted by spotting cultures onto 0.3% (wt/vol) FAB soft agar. The absence of a halo around the initial inoculum indicates loss of flagellum-driven motility. Isolates with flagellar mutations (*fleQ* and *flgA*) showed reduced swimming motility relative to the ancestor. Isolates with the *gacA* mutation and enhanced swimming phenotype (mSwim) exhibited increased swimming motility relative to the ancestor. Isolates with the *ΔcheA* mutation exhibited swimming motility comparable to the ancestral population.

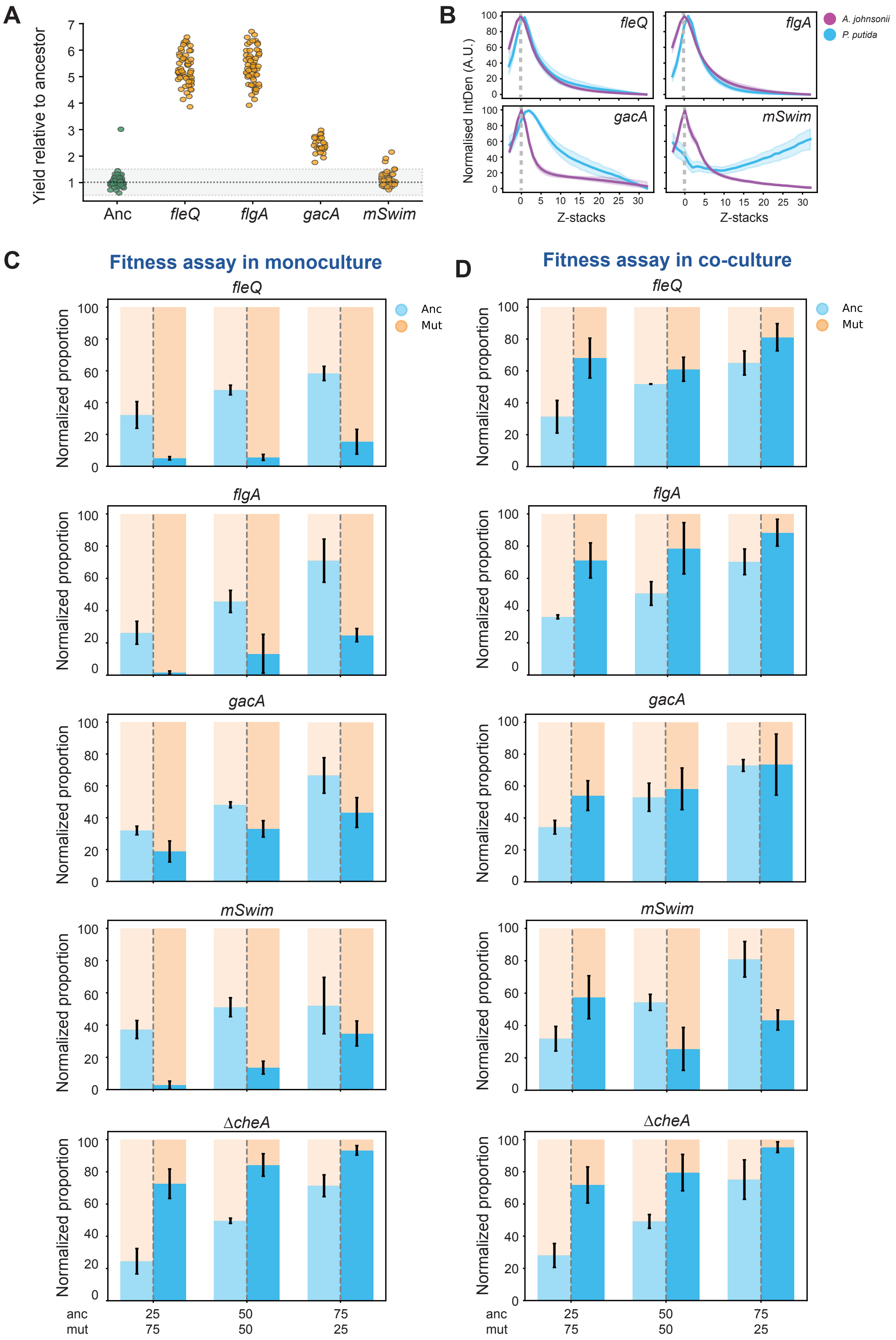

A
Yield relative to ancestor
Anc
fleQ
flgA
gacA
mSwim
B
Normalised IntDen (A.U.)
A. johnsonii
P. putida
Z-stacks
C
Fitness assay in monoculture
D
Fitness assay in co-culture
Normalized proportion
Anc
Mut
ΔcheA
anc
mut

**Figure S5.** (*A*) Scatterplot showing the yield relative to the ancestral population for 60 colonies derived from clonal populations carrying specific mutations (*fleQ, flegA, gacA* and mSwim).The gray area represents the zone of non-significant differences. (*B*) Normalized integrated density fluorescent signal corresponding to *A. johnsonii* (magenta), and a clonal population of *P. putida* (cyan) carrying specific mutations (*fleQ, flegA, gacA* and mSwim) across the z range (from 0 to 30 µm above the bottom of the well with an image taken with a 1 µm increment). Dark-colored lines correspond to the median values of 3 different fields of view per lineage tested, and light-colored areas correspond to the 95% CI. (*C-D*) Histograms showing the normalized proportions of ancestral (blue) and mutant (orange) populations at t0 (left of the dashed gray line) and t24 (right of the dashed gray line) across different initial ratios in competition fitness assay between the ancestral and evolved *P. putida* populations in monoculture (panel *C*) and in co-culture with *A. johnsonii* (panel *D*).

#### Supplementary Tables

**Table S1.** Representative mutations well-established in evolved populations.

| gene | mutation | annotation | description |
|---|---|---|---|
| *fleQ* | Δ13 bp | coding (1241-1253/1476 nt) | transcriptional regulator FleQ |
| *flgA* | C→T | Q184* (CAA→TAA) | flagellar basal body P-ring formation chaperone FlgA |
| *flgH* | +C | coding (38/696 nt) | flagellar basal body L-ring protein FlgH |
| *flhA* | Δ1 bp | coding (966/2130 nt) | flagellar biosynthesis protein FlhA |
| *gacA* | T→G | H73P (CAC→CCC) | UvrY/SirA/GacA family response regulator transcription factor |
| *gacS* | (GGTGGCCAG GTCGCGGT)$_{1\rightarrow2}$ | coding (401/2754 nt) | GacS response regulator |

**Table S2.** All mutations identified in populations using *breseq*.

A separate file (.xlsx) is provided

**Table S3.** List of strains, plasmids, and primers used in this study.

| Strain ID | Drug resistance | Species | Genotype/Background |
|---|---|---|---|
| TBRO3 | Amp$^r$/Gm$^r$ | *E. coli* | *ΔcheA* + *kan* cassette in modified PTW475 plasmid |
| TBRO4 | Kan$^r$/ Gm$^r$ | *P. putida* | *ΔcheA* + *kan* + *gfp* |
| TBRO7 | Gm$^r$ | *P. putida* | *gfp;* ancestor |
| TBRO8 | Strep$^r$ | *A. johnsonii* | ancestor |
| TBRO9 | Strep$^r$ / Kan$^r$ | *A. johnsonii* | *rfp* |
| TBRO11 | Gm$^r$ | *P. putida* | *gacA* mutation (T→G) in 200g evolved obligate cross-feeding lineage 4 |

| | | | |
|---|---|---|---|
| TBRO16 | Gm$^r$ | *P. putida* | *fleQ* mutation (Δ13) in 200g evolved non-biogenic low lineage 4 |
| TBRO17 | Gm$^r$ | *P. putida* | *flgA* mutation (C→T) in 200g evolved non-biogenic low lineage 6 |
| TBRO19 | Gm$^r$ | *P. putida* | mSwim in 200g evolved obligate cross-feeding lineage 6 |
| **Plasmid ID** | **Drug resistance** | **Background vector** | **Insert fragment** |
| pTBRO1 | Amp$^r$ / Gm$^r$ | pTW475 | - |
| pTBRO3 | Amp$^r$ / Gm$^r$ / Kan$^r$ | pTW475 | *ΔcheA* + *kan* cassette (flanking upstream and downstream region of *cheA* gene followed by *kan* resistance gene) |
| **Primer Name** | **Sequence** | | |
| gacA_f | TTTTACAGGCTTGCGTCAACCAT | | |
| gacA_r | TGTGCTTGATTAGGGTCTTAGTGGTC | | |
| fleQ_f | GGGGGTTTGGGTTTGACGTA | | |
| fleQ_r | GACGTCCGAACACTGACCAT | | |
| flgA_f | CGTTTCAGCCTTTCGTTGCA | | |

| | |
|---|---|
| flgA_r | CGCTGATCTTCTGCAACTGC |
| fleQ_f_2 | ACACCCAGCCCAAACATCAA |
| fleQ_r_2 | ATCCTTGGTCGCACGAATGT |
| flgA_f_2 | AGGGCATTGCAGTACTTCCC |
| flgA_r_2 | TTGCCTTGTTTTCGGCCAAC |
| pTW475_Insert-check_F | AGGCATCAAATAAAACGAAAGGCT |
| pTW475_Insert-check_R | GCCAGCTAGAGGACCAGC |
| cheA_FC_f | GCGAGTTTAAACCCCCTGGCATAGCTGCCG |
| cheA_FC_r_3 | CAAAATACCTCGGCGTATTTGATTTCGG |
| cheA_RC_f_3 | GCGGGACTCTGAAGCTCATCAAACGTGC |
| cheA_RC_r | AGTCTGGTCGACCCCGCACCATGGACCTGG |
| Kan_for_gibson_2 | AAATACGCCGGTTTTATGGACAGCAAGCGAA |
| Kan_rev_gibson_2 | GATGAGCTTCAGAGTCCCGCTCAGAAGAACT |

#### Movie legends

Link to Movie S1.

**Movie S1.** Video of a representative 3D image of the co-culture assay showing ancestral *P. putida* (cyan) with ancestral *A. johnsonii* (magenta) at 72 h post-inoculation. The movie shows the Z-stacks moving from the bottom of the well (5 μm below the glass-media interface) to the interior of the well (30 μm above the glass surface).

Link to Movie S2.

**Movie S2**. Video of a representative 3D image of the co-culture assay showing the lineages evolved under non-biogenic high conditions of *P. putida* (cyan) with ancestral *A. johnsonii* (magenta) at 72 h post-inoculation. The movie shows the Z-stacks moving from the bottom of the well (5 μm below the glass-media interface) to the interior of the well (30 μm above the glass surface).

Link to Movie S3.

**Movie S3**. Video of a representative 3D image of the co-culture assay showing the lineages evolved under non-biogenic low conditions of *P. putida* (cyan) with ancestral *A. johnsonii* (magenta) at 72 h post-inoculation. The movie shows the Z-stacks moving from the bottom of the well (5 μm below the glass-media interface) to the interior of the well (30 μm above the glass surface).

Link to Movie S4.

**Movie S4.** Video of a representative 3D image of the co-culture assay showing the lineages evolved under facultative cross-feeding conditions of *P. putida* (cyan) with ancestral *A. johnsonii* (magenta) at 72 h post-inoculation. The movie shows the Z-stacks moving from the bottom of the well (5 µm below the glass-media interface) to the interior of the well (30 µm above the glass surface).

Link to Movie S5.

**Movie S5.** Video of a representative 3D image of the co-culture assay showing lineages evolved under obligate cross-feeding conditions of *P. putida* (cyan) with ancestral *A. johnsonii* (magenta) at 72 h post-inoculation. The movie shows the Z-stacks moving from the bottom of the well (5 µm below the glass-media interface) to the interior of the well (30 µm above the glass surface).

Link to Movie S6.

**Movie S6.** Representative time-lapse video of the co-culture microscopic assay for the ancestral *P. putida* (cyan) with ancestral *A. johnsonii* (magenta), 15 µm above the glass surface.

Link to Movie S7.

**Movie S7.** Representative time-lapse video of the co-culture microscopic assay for the lineages evolved under obligate cross-feeding conditions of *P. putida* (cyan) with ancestral *A. johnsonii* (magenta), 15 µm above the glass surface.
